# Transmembrane Signaling Regulates the Remodeling of the Actin Cytoskeleton: Roles of PKC and Adducin

**DOI:** 10.1101/2021.09.29.462380

**Authors:** Bih-Hwa Shieh, Wesley Sun, Darwin Ferng

## Abstract

We tested the hypothesis that Pkc53E regulates adducin to orchestrate the remodeling of the membrane skeleton following the transmembrane GPCR-Gq signaling. Adducin is a known substrate of PKC and is critical for the assembly of the membrane skeleton by cross-linking actin filaments with the spectrin network. In *Drosophila* photoreceptors, loss of function in *pkc53E* leads to retinal degeneration while Pkc53E-RNAi negatively impacts the actin cytoskeleton of the visual organelle rhabdomeres. Unexpectedly, Pkc53E-RNAi enhances the degeneration caused by the loss of PLCβ4 (*norpA^P24^*). We show that when PLCβ4 is absent Plc21C may be activated instead for activating Pkc53E. We investigate whether Pkc53E phosphorylates adducin *in vivo* and observed that levels of phosphorylated adducin at the conserved PKC site were greatly reduced in a null allele of *pkc53E*. We show Pkc53E-RNAi did not modify adducin-RNAi, which exerts a more severe effect on the actin cytoskeleton. Moreover, overexpression of the mCherry-tagged adducin that appears to act in a dominant-negative manner interferes with the spectrin interaction leading to the apical expansion of rhabdomeres similar to that of β-spectrin-RNAi. We performed epistasis analysis and show that double mutants of the tagged adducin and Pkc53E-RNAi display the expansion phenotype at the eclosion, but progress to severe degeneration in adult photoreceptors. Together, most of our findings support that adducin is likely regulated by Pkc53E in *Drosophila* photoreceptors.

## Introduction

Conventional protein kinase C (cPKC) is a target of phorbol esters and is activated by diacylglycerol (DAG) and Ca^2+^ (Lipp and Reither, 2011). cPKC is a downstream mediator following the activation of phospholipase C (PLC) including PLCβ and PLCγ. PLCγ is activated following the stimulation of growth factor receptors, and cPKC has been shown critical for processes associated with morphological changes leading to growth and differentiation of cells. In contrast, in PLCβ mediated signaling events such as the visual signaling that takes place in *Drosophila* photoreceptors, in which rhodopsin couples to the heterotrimeric Gq protein leading to the activation of PLCβ4 (NorpA) (Hardie and Juusola, 2015; Montell, 2012), the role of cPKC is less explored. cPKC has been commonly linked to the regulation of the actin cytoskeleton (Larsson, 2006). Indeed, several PKC substrates have been identified and characterized. However, the mechanisms by which cPKC regulates cell morphology or the cytoskeleton remain elusive.

There are two cPKCs expressed in *Drosophila* photoreceptors, eye-PKC (Schaeffer et al., 1989) and Pkc53E (Rosenthal et al., 1987), both of which share over 70% sequence identities. Eye-PKC is expressed in photoreceptors and is localized in the rhabdomere (Smith et al., 1991), the visual organelle in which the visual signaling takes place. Moreover, eye-PKC is an integral part of the multimeric signaling complex organized by the scaffolding protein INAD (inactivation-no-afterpotential D) (Adamski et al., 1998; Tsunoda and Zuker, 1999). Eye-PKC has been shown to phosphorylate both INAD and TRP (transient receptor potential) (Peng et al., 2008; Popescu et al., 2006). In contrast, the role of Pkc53E has not been investigated.

Studies have linked cPKC to the regulation of the actin cytoskeleton (Larsson, 2006). The actin cytoskeleton consists of a network of actin microfilaments, which can be found in the cell cortex, the stress fiber, and various extensions including filopodia, lamellipodia, and microvilli. The actin cytoskeleton is critical for maintaining cell shape and regulates diverse processes including endocytosis, cytokinesis, and chemotaxis. In the cell cortex, actin microfilaments support the plasma membrane by forming the membrane scaffold with the spectrin network (Baines, 2010). Moreover, the interaction between spectrin and actin filaments is dynamically regulated by adducin (Baines, 2010), a family of cytoskeletal proteins consisting of three adducins (α-, β- and γ-adducin) (Joshi et al., 1991). Importantly, adducin is regulated by Ca^2+^ and by PKC as adducin contains the sequence related to the conserved MARCKS (myristoylated alanine-rich C-kinase substrate) domain. For example, in platelets activation by par ligands or phorbol esters results in PKC phosphorylation of adducin, which leads to its dissociation from spectrin and actin (Barkalow et al., 2003). The release of phosphorylated adducin from the barbed end of the actin filament would promote actin polymerization. Thus, adducin acts as a key regulatory protein to initiate remodeling of the actin cytoskeleton following an increase of the intracellular Ca^2+^ and activation of PKC. Pkc53E may regulate adducin to orchestrate reorganization of the actin cytoskeleton following the activation of the visual signaling.

Here we characterized a photoreceptor-specific isoform of Pkc53E, Pkc53E-B, and show that downregulation leads to a loss of rhabdomeres possibly by affecting the actin cytoskeleton. We explored its regulation by the visual signaling and observed Pkc53E was partially active in the absence of PLCβ4 /NorpA. Indeed, we show Plc21C (Shortridge et al., 1991) may be involved when NorpA is absent. We investigated whether Pkc53E regulates *Drosophila* adducin or Hts (hu-li tai shao) (Yue and Spradling, 1992) and demonstrate that phosphorylation of adducin at the conserved PKC site was greatly reduced in *pkc53E* mutants. Moreover, downregulation of adducin by RNAi results in loss of rhabdomeres. In contrast, overexpression of a tagged adducin isoform leads to apical expansion of rhabdomeres similar to the knockdown of spectrins. Epistasis analysis supports the notion that Pkc53E is upstream of adducin during rhabdomere morphogenesis. Furthermore, the dominant-active adducin potentiates the deterioration of the actin cytoskeleton triggered by Pkc53E-RNAi.

## Materials and Methods

### Fluorescence microscopy

Adult flies were anesthetized by CO_2_, immobilized in clay with compound eyes facing upward in a 50 mm Petri dish for imaging. The compound eye was examined using an upright Olympus AX70 microscope equipped with a 10X lens (for detecting deep pseudopupil) or 40X (LUMPlan 40X) or 100X (LUMPlan 100X) water immersion lens for examining multiple ommatidia. Image acquisition was performed at 100X magnification for dpp and 400X or 1000X for rhabdomeres/ommatidia using IPLab image acquisition software (BioVision Technologies, Exton, PA, USA) and the Retiga camera from QImaging (Surrey, BC, Canada). Exposure time was made constant throughout each experiment based on the brightest signal in the control group. Multiple flies (n ≥ 3) of each genotype were analyzed.

### Fly handling for microscopy

Adult flies were sorted and manipulated under a dissecting microscope with a light source of 600 lux for less than one minute. During imaging, compound eyes were subjected to the blue light (1300 lux) or green light (4500 lux) from the fluorescent microscope, and images were taken immediately for Actin-GFP, Arr2-GFP, or Rh1-mCherry.

### Fluorescent image analysis

All image manipulation was performed under the guideline of Rossner and Yamada (Rossner and Yamada, 2004). Fluorescent images included in the Figures are similar in appearance to the raw images. Experimentally, we collected newly eclosed flies of the desirable genotype, placed them in a vial, and aged at 25 °C in a 12h light/dark cycle (ambient light 300 lux) for various amounts of time. Flies were analyzed for GFP-marked rhabdomeres by water-immersion fluorescence microscopy. Retinal morphology was scored based on the number of rhabdomeres present in a given ommatidium (unit eye) similar to that described by Cerny et al. (Cerny et al., 2013). Briefly, retinas from three flies from each eight ommatidia were counted to obtain the average of the rhabdomere number. To score rhabdomere areas, ImageJ was used. Four ommatidia clusters were selected in which the total rhabdomere area of each cluster was measured and averaged. Wild-type flies of the same age were used as controls.

### Quantitative Western blotting

Single fly head or four retinas was dissected and extracted with 15 μl of 2X Laemmli sample buffer by sonication. Protein extracts were size-fractionated by SDS/PAGE (10-12%) and transferred onto a nitrocellulose filter. Filters were incubated with desired primary antibodies, followed by the fluorophore-conjugated secondary antibodies (IRDye 680LT Goat anti-Rabbit IgG, or IRDye 800CW Goat anti-mouse IgG, LI-COR). The fluorophore signal was detected by the Odyssey Infrared Imaging System (LI-COR, Lincoln, NE, USA), and analyzed by Image Studio™ 5.2.

Individual protein content was normalized by comparing it to the loading control (actin or fascin). We carried out three to five analyses (n = 3-5) with each using one fly head or four retinas. Polyclonal antibodies against Gq α-subunit were generated in rabbits using a bacterial fusion protein corresponding to 1-200 aa of Gq. Polyclonal anti-phospho-Ser^726/705/703^ adducin antibodies were purchased from Millipore (Burlington, MA, USA). Monoclonal antibodies for Rh1 (4C5), TRP (MAb83F6), adducin/Hts (1B1), actin (JLA20), and fascin (sn_7C) were obtained from Developmental Studies Hybridoma Bank (University of Iowa).

#### Recombinant DNA and Molecular Biology

A full-length Rh1 cDNA without the stop codon was generated by PCR with the engineered SacI (5’) and EcoRI (3’) restriction enzyme sites. The mCherry cDNA sequence with the flanking EcoRI (5’) and XhoI (3’) restriction enzyme sites was generated by PCR using pUAST-mCherry [a gift from Dr. Amy Kiger (UCSD)] as the template. The mCherry nucleotide sequence was inserted in-frame into the 3’ of the Rh1 cDNA and the resulting Rh1-mCherry chimera DNA was subcloned into YC4 for the expression under the control of the *Drosophila* Rh1 promoter (Kristaponyte et al., 2012).

### Reverse Transcription-Polymerase Chain Reaction (RT-PCR)

Total RNA was extracted from 20 fly heads (or 50 retinas) by a modified method of Chirqwin et al (Chirgwin et al., 1979) and dissolved in 20 μl water. Five microliter (μl) of total RNA were used for the first-strand cDNA synthesis via Superscript III (Invitrogen, Carlsbad, CA) primed with random hexamers. Quantitative PCR in triplicate was performed via CFX96 Real-Time System (BIO-RAD, Hercules, CA) using iQTM SYBR Green Supermix (BIORAD). All expression values were normalized to RpL32 (rp49). To compare isoform specific transcript in various genetic backgrounds, RT/PCR products were analyzed by 8% polyacrylamide gel and relative band intensity was quantified using Image Lab (Bio-Rad). Most of the primer sequences used were selected from the FlyPrimerBank (Hu et al., 2013) and listed below: *rp49* (113 nt), AGCATACAGGCCCAAGATCG (5’), TGTTGTCGATACCCTTGGGC (3’); *Pkc53E* (for total *Pkc53E*, 90nt), AGACTCGCACCATTAAGGCTT (5’), GGATGCGTCGATCCTTGTCTT (3’); *Pkc53E* (for distinguishing between C/E/B, 356/359 nt and A/F, 332 nt), CACGTTCTGCTCCCACTGCA (5’), GCTCCGTGTGATCGCATCC (3’); *Pkc53E* (for F, 262 nt or B, 289 nt), AGCCCTCAAGAAGAAGAACGT (5’), TCCTGCGTATGTGAATGGCTC (3’); *Pkc53E* for C/E (282 nt) AGCCCTCAAGAAGAAGAACGT (5’), AGGAAGGTGACATTCTGCCA (3’); *α-spectrin* (205 nt), GGTTCTGTCCCGCTACAATG (5’), TCGAACGCCTGATGTTTCTGG (3’); *β-spectrin* (121 nt) CCGCATTGCCGATCTCTATGT (5’), CAGGCAGTGGATACGCATCTT (3’); *karst* (182 nt), GTCGGAGAAACTGGGCAAG (5’), GAAACCGCAGAATGATGGTCC (3’); *arr1* (90 nt), CATGAACAGGCGTGATTTTGTAG (5’), TTCTGGCGCACGTACTCATC (3’); *Plc21C* (159 nt), GAGAAGACAGTGACGGTATGC (5’), CAGGAACATAATCGCCGAGC (3’); Pld (132 nt), GATGAGACCCTCGCTTTTCCT (5’), GACTACACTGTTGTTTTCCTCGT(3’). Primers specifically for isoforms I/L/M of adducin are AGAGCAGTAAAGAGTTCCAG (5’) and GGCCTCGGCCTTCTTCTTC(3’), for isoforms O/S are CAGAGCAGTAAAGAGGATGTC (5’) and GGCCTCGGCCTTCTTCTTC(3’). Primers for distinguishing between Q/R or O/S are CCACAAAGCGAACCCGAGA (5’) and GACATCCTCTTTACTGCTCTG (3’).

### Drosophila stocks

*Drosophila* lines including mutants for eye-PKC (*inaC^P209^*, #42241), and *Pkc53E* (*Pkc53ED^Δ28^*, #80988) were obtained from Bloomington Drosophila Stock Center (BDSC) (NIH P40OD018537). Transgenic flies for GMR-GAL4 (stock #1104), RNAi lines for the following genes including *hts* (#35421), *α-spectrin* (#48201), *β-spectrin* (#48202), *spectrin-βH* or *karst* (#33933), *inaC* (#36776), *pkc53E* (#55864), *Gaq* (#63987), *plc21C(#31269*, #33719), and *pld* (#32839) were also obtained from BDSC. UAS-driven overexpressing lines including *hts-mCherry* (#66171), *Pkc53E-B* (#80989), and Actin-GFP (#9253) were from BDSC. Fly cultures were maintained in the standard cornmeal medium at 25 °C. Standard crosses were used to introduce suitable genetic backgrounds.

### Statistical analysis

One-way ANOVA and two-tailed Student’s *t*-test were employed for statistical analysis.

## Results

### Characterization of a *pkc53E* null allele and identification of a photoreceptor-specific Pkc53E isoform

In *Drosophila* photoreceptors, activation of rhodopsin leads to a transient rise of cytoplasmic Ca^2+^, which has been implicated in regulating the visual response and in maintaining the structural integrity of photoreceptors (Voolstra and Huber, 2020). The Ca^2+^ transient thereby activates protein kinases including the Ca^2+^/calmodulin-dependent protein kinase II and PKC. In the *Drosophila* genome, there are two cPKCs, eye-PKC and Pkc53E, both of which are expressed in the eye. Specifically, eye-PKC has been shown to phosphorylate and regulate TRP (Peng et al., 2008; Popescu et al., 2006; Voolstra et al., 2013), and mutants lacking eye-PKC (*inaC*) display abnormal visual signaling and retinal degeneration (Hardie et al., 1993; Ranganathan et al., 1991; Smith et al., 1991). In contrast, the role of Pkc53E remains elusive. We explored the expression and function of Pkc53E in photoreceptors.

First, we characterized a null allele of *pkc53E, pkc53E^Δ28^* (Figure S1), and show it displays light-dependent retinal degeneration that is accompanied by a reduction of Rh1(Figure 1A). The loss of Rh1 is less severe compared to a null allele of eye-PKC (*inaC^P209^*) as seven-day-old *pkc53E^Δ28^* contained about 47.8±6.1% (n = 4) of Rh1, while *inaC^P209^*, 14.8±2.5% (n = 4) by Western blotting.

**Figure 1:**
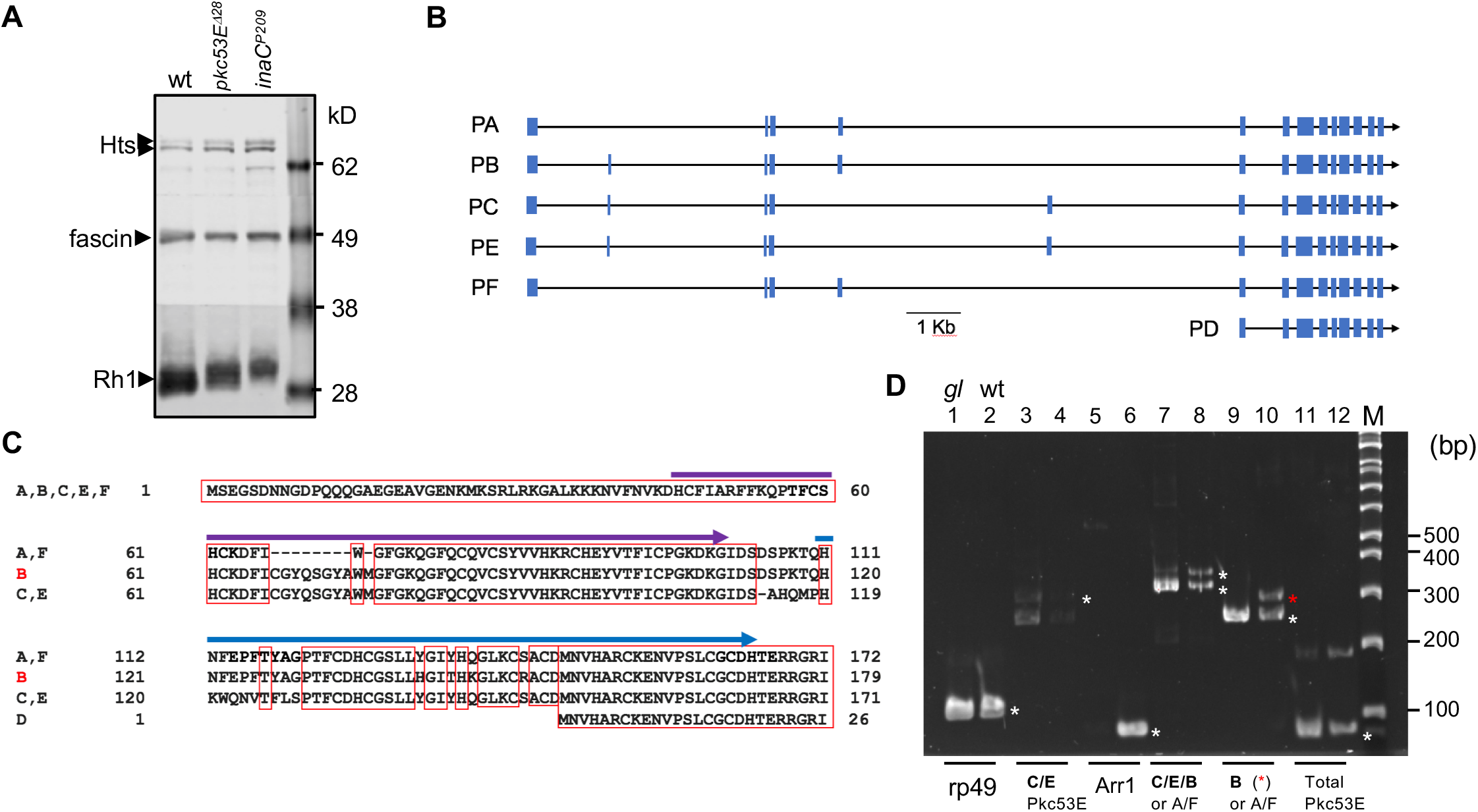
Characterization of a *pkc53E* null allele and identification of a photoreceptor-specific Pkc53E isoform. **A**, Loss of function in *pkc53E* leads to retinal degeneration. Shown is a Western blot containing extracts from wildtype, *pkc53ED^Δ28^*, and *inaC^P209^* (seven-day-old) probed with antibodies against Rh1, fascin (loading control), and adducin/Hts, respectively. Protein standards are indicated on the right. **B**, A graphic map depicting the coding sequences (filled boxes) of six alternatively spliced *pkc53E* transcripts, A-F. **C**, Alignment of the N-terminal sequences from the six Pkc53E isoforms. All isoforms have two distinct C1/DAG binding domains (arrows, 45-110 aa; 120-173 aa in isoform B), except the D isoform. Amino acid identities are boxed. **D**, Identification of the photoreceptor-specific isoform by RT/PCR. Shown are PCR products analyzed by polyacrylamide gel (8%). Even-numbered lanes represent PCR products from wild-type and odd-numbered lanes, *glass* mutants. Rp49 serves as a positive control whereas Arr1, a positive control for wild-type and negative control for *glass* for its expression in photoreceptors. DNA fragments corresponding to the predicted PCR products are marked with asterisks (*) in wild-type lanes. The B isoform of Pkc53E appears highly expressed in photoreceptors as it is drastically reduced in the *glass* mutant (lanes 9). DNA size standards are shown on the right.

Next, we investigated the critical Pkc53E isoform in photoreceptors. The *pkc53E* gene encodes six transcripts (RA-F) leading to the translation of four distinct polypeptides, which include isoforms of 679 amino acids (aa) (PB), 678 aa (PC, PE), 670 aa (PA, PF), and 525 aa (PD), with different N-terminal sequences (Figure 1B). Specifically, all isoforms contain the C2 domain for the Ca^2+^ binding and all also contain two C1 domains that bind to DAG, except PD (Figure 1C). PD lacks the N-terminal 145 aa and therefore its activity is likely independent of DAG. DAG may exert different affinity towards PA and PF isoforms as both are missing eight amino acids in the first C1 domain when compared to the PB, PC, and PE isoforms (Figure 1C). Taken together, these Pkc53E isoforms have distinct C1 domains, which regulate the activation of each isoform.

We investigated which *pkc53E* isoforms are expressed in photoreceptors via RT/PCR (Figure 1D). Using oligonucleotide primers flanking or at the alternative spliced sites, we demonstrate that mRNA coding for the isoform PB/PC/PE group is present in wild-type (Figure 1D, lane 8) but greatly reduced in heads of *glass* mutants lacking photoreceptors (Moses et al., 1989) (lane 7). This result supports that the group of the PB/PC/PE transcripts is preferentially expressed in photoreceptors. To explore further, we observed that transcripts for the PC/PE group are present at a low level in both wild-type and *glass* heads (Figure 1D, lanes 3 and 4). In contrast, mRNA for the PB isoform is highly expressed in wild-type (Figure 1D, lane 10) but not *glass* heads (lane 9). Taken together, our data support *pkc53E-B* that encodes a protein of 679 aa is preferentially expressed in photoreceptors.

### Downregulation of either Pkc53E or eye-PKC leads to retinal degeneration via two GFP reporters

To investigate the role of Pkc53E-B in photoreceptors, we employed RNAi-mediated strategies (Perrimon et al., 2010) using the GMR driver that targets photoreceptors (Freeman, 1996). First, we validated RNAi by RT/PCR and observed that mRNA corresponding to Pkc53E-B is greatly reduced to about 19 % of wild-type [from 45.3 ± 8.5 (n = 3) to 8.9 ± 5.1 (n = 3)] (Lanes 5, 9, Figure 2A). We also performed eye-PKC-RNAi, which was validated by Western blotting with a great reduction of eye-PKC to 4.9 ± 3.4% (n = 4) of wild-type (Figure S1). Subsequently, we investigated the retinal morphology by fluorescence microscopy using GFP tagged arrestin 2 (Arr2-GFP) expressed in R1-6 photoreceptors (Kristaponyte et al., 2012) as the reporter. Arr2 binds to activated rhodopsin in the rhabdomere.

**Figure 2:**
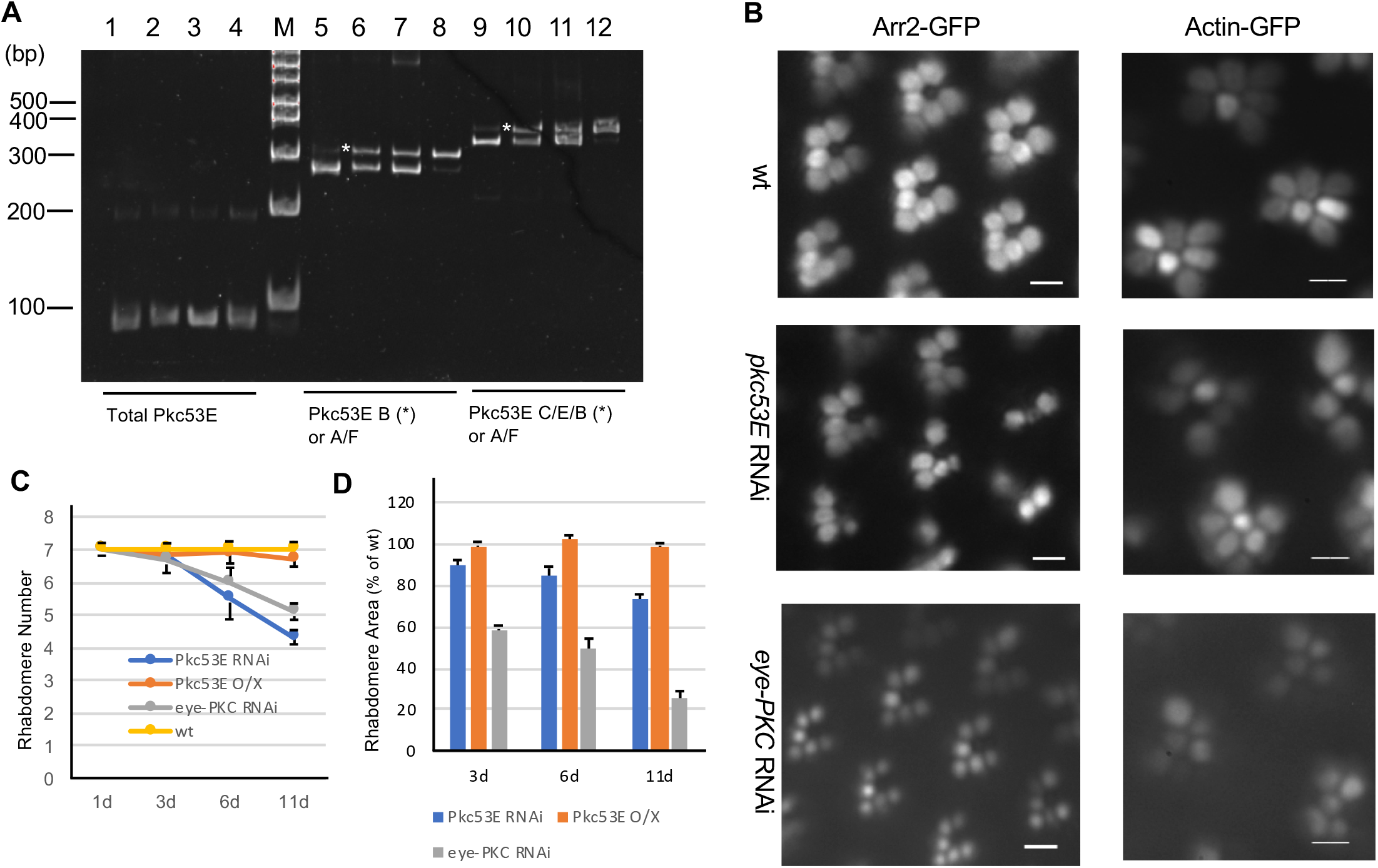
Downregulation of Pkc53E or eye-PKC in photoreceptors leads to distinct degeneration phenotype. **A**, Validation of RNAi and overexpression of *pkc53E* by RT/PCR. The mRNA level for the B-isoform of Pkc53E (*) is drastically reduced to 8.9 ± 5.1 (n = 3) when compared to the control of 45.3 ± 8.5 (n = 3, lanes 5, 9). In contrast, overexpression of *pkc53E-B* leads to about a two-fold increase of the mRNA (lanes 8, 12). Oligonucleotide primer sets used are indicated below and the first-strand cDNA templates from Pkc53E-RNAi (lanes 1, 5, and 9), wild-type (2, 6, and 10), flies expressing GMR driver alone (3, 7, and 11), and *pkc53E* overexpressing flies (4, 8, and 12), respectively, were used. The PCR products corresponding to the B-isoform are marked with asterisks on the left in the wild-type lanes. **B**, Retinal degeneration caused by Pkc53E-RNAi or eye-PKC-RNAi using Arr2-GFP (left) or Actin-GFP (right) as the reporter. Arr2-GFP was engineered to be expressed in R1-6 photoreceptors whereas Actin-GFP was targeted to all photoreceptors (R1-R8) driven by the GMR driver. Shown are representative images of 8-day old flies. Scale bars on the left panel, 5 μm, and on the right panel, 2 μm. **C**, The age-dependent loss of rhabdomeres in the *pkc* (Pkc53E RNAi, Pkc53E overexpression, eye-PKC RNAi) mutants using Actin-GFP as the reporter. Each time point represents the mean of three flies (mean ± S.E.M, n = 3). **D**, The age-dependent changes of rhabdomere area in various *pkc* mutant backgrounds. Shown are mean ± S.E.M (n = 3) from three independent experiments.

We show eye-PKC-RNAi led to more severe degeneration affecting mostly size initially with less impact on the number, orientation, and arrangement of rhabdomeres within the cluster (Figure 2B). In contrast, Pkc53E-RNAi resulted in an age-dependent retinal degeneration characterized by distorted ommatidia clusters with missing rhabdomeres (Figure 2B). Indeed, in 8-day old retinas, about one to two rhabdomeres were missing in most ommatidia clusters with 4.8 ± 0.7 (n = 3) remaining. In contrast, overexpression of Pkc53E-B, which increases the mRNA content about two-fold (203.5 ±17.2%, n = 3) (Figure 2A), did not significantly modify the retinal morphology (Figure 2C, D; Figure 8A). Together, our data support that both Pkc53E and eye-PKC are involved in the regulation of the rhabdomere integrity as reducing the expression of each kinase leads to distinct degeneration phenotypes.

Conventional PKC has been implicated in regulating the actin cytoskeleton. Therefore, we examined how the actin-based rhabdomeres might be affected using Actin-GFP that is expressed in R1-8 photoreceptors as the reporter (Roper et al., 2005; Shieh et al., 2021). Indeed, Pkc53E-RNAi results in missing rhabdomeres, which is accompanied by the abnormal arrangement of ommatidia clusters (Figure 2B). We show that the number of rhabdomeres is reduced from seven to about five in 10-day old retinas, which is similar to that observed using Arr2-GFP. In contrast, eye-PKC-RNAi initially affects the rhabdomere diameter, but not the rhabdomere number (Figure 2B). The age-dependent reduction of the rhabdomere number (Figure 2C) or the area (Figure 2D) upon knockdown or overexpression of Pkc53E was compared.

Together, our findings strongly support that a reduction of the Pkc53E activity directly or indirectly affects the actin cytoskeleton of rhabdomeres. We thus propose that Pkc53E may regulate processes to promote the stability of rhabdomeres.

### Downregulation of either Pkc53E or eye-PKC by RNAi enhanced the degeneration of *norpA^P24^* photoreceptors

We explored whether Pkc53E is operating in the visual cascade downstream of NorpA, similar to eye-PKC. We speculate that both eye-PKC and Pkc53E would be inactive in *norpA^P24^* photoreceptors lacking NorpA/PLCβ4. Consequently, degeneration of *norpA^P24^* photoreceptors would not be modified by downregulation of each PKC. Unexpectedly, we observed that Pkc53E-RNAi greatly exacerbated retinal degeneration of *norpA^P24^* photoreceptors (Figure 3A, D). Similarly, eye-PKC-RNAi also exerted a similar but less severe effect (Figure 3A, D). Taken together, we propose that both eye-PKC and Pkc53E remain active in mutants missing PLCβ4, suggesting that activation of either another PLCβ or alternate pathways leading to the synthesis of DAG, such as the activation of phospholipase D (Pld) (LaLonde et al., 2005) may be involved.

**Figure 3:**
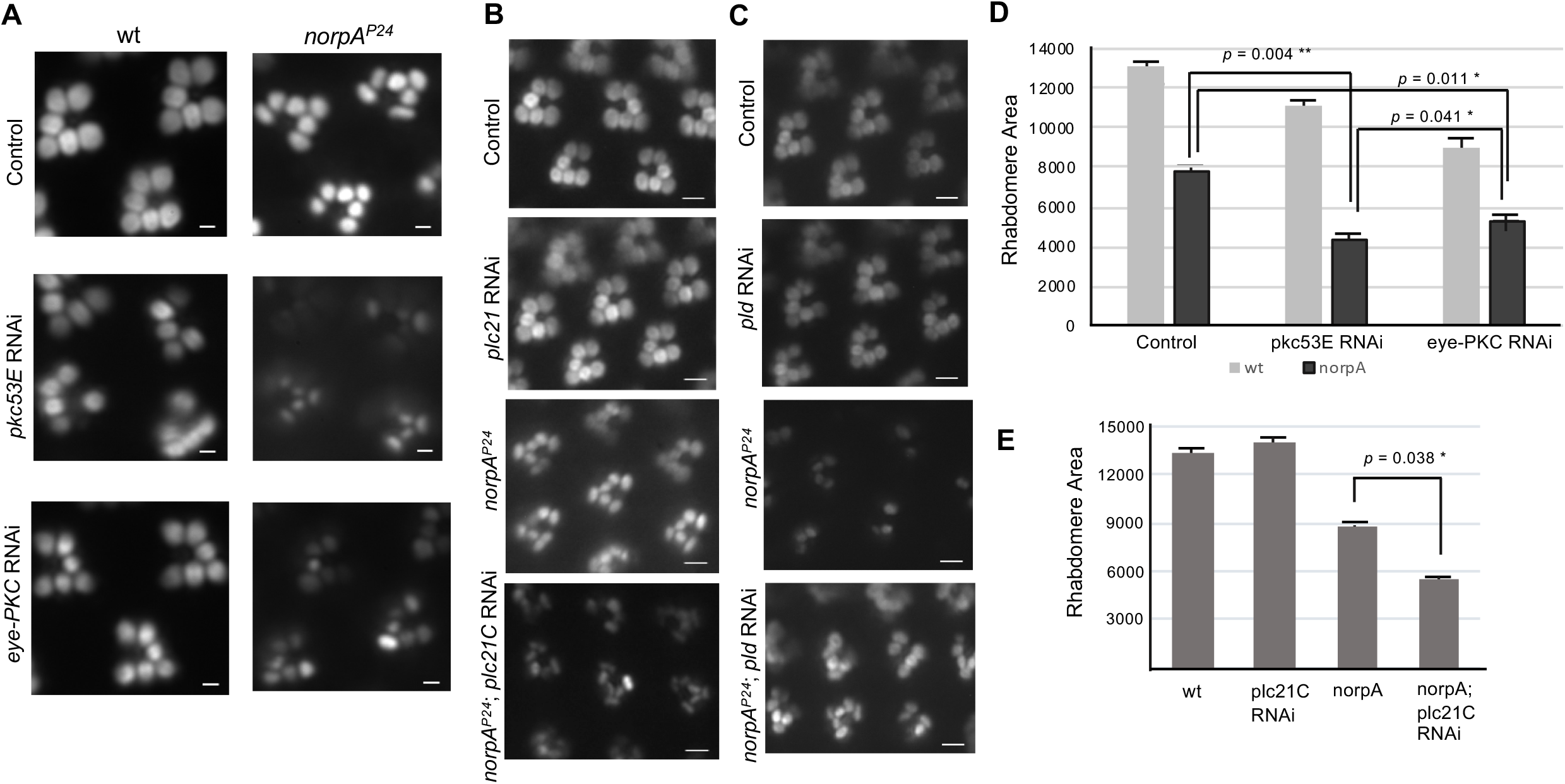
Activation of Plc21C may be activated when NorpA/PLCβ4 is absent. **A**, Retinal degeneration of *norpA^P24^* is enhanced by either Pkc53E-RNAi or eye-PKC-RNAi. Shown are retinal morphology in 7-day old flies in either wild-type (left panel) or *norpA^P24^* (right) genetic background. Arr2-GFP was used as the reporter. Scale bars, 2 μm. **B**, Degeneration of *norpA^P24^* photoreceptors is enhanced by Plc21C-RNAi. In contrast, Plc21C-RNAi alone did not affect the retinal morphology of wild-type. Shown are retinas of 7-day old flies. Scale bars, 5 μm. **C**, Degeneration of *norpA^P24^* photoreceptors is delayed by Pld-RNAi. In contrast, Pld-RNAi alone did not modify the retinal morphology of wild-type photoreceptors. Shown are retinas of 10-day old flies. Scale bars, 5 μm. **D**, **E**, Comparison of rhabdomere areas (in arbitrary units, n = 4) in various genetic backgrounds.

### Plc21C-RNAi but not Pld-RNAi enhanced the degeneration of *norpA^P24^* photoreceptors

We investigated the contribution of phospholipase C at 21C (Plc21C), which is known to couple to Gq in *norpA^P24^* photoreceptors (Ogueta et al., 2018). Plc21C functions in olfaction (Kain et al., 2008) and also has been shown to participate in the light-dependent regulation of the circadian clock (Ogueta et al., 2018). We speculate if Plc21C promotes the activation of two cPKC’s, downregulation of Plc21C would also exacerbate retinal degeneration of *norpA^P24^* photoreceptors. Indeed, we observed that retinal degeneration was greatly accelerated by Plc21C-RNAi, as evident in 7-day old flies (Figure 3B, E). However, Plc21C-RNAi did not modify the retinal morphology of wild-photoreceptors (Figure 3B, E), indicating that it is not involved in the maintenance of rhabdomeres. In contrast, Pld-RNAi delayed the *norpA^P24^* degeneration, while it did not modify wild-type retinal morphology (Figure 3C). Quantitative PCR analyses indicated that mRNA for Pld and Plc21C in the fly head was reduced by 55 ± 14% (n = 3), and 41 ± 11 % (n = 3), respectively, following RNAi-mediated downregulation.

Taken together, our findings suggest the contribution of Plc21C but not Pld leading to the generation of DAG thereby activating Pkc53E and eye-PKC in the absence of NorpA/PLCβ4.

### Gqα-RNAi leads to retinal degeneration, which is not modified by Pkc53E-RNAi

We explored how Plc21C might be activated in *norpA^P24^* photoreceptors. Like PLCβ4, activation of Plc21C also involves the heterotrimeric Gq protein (Kain et al., 2008). We investigated the role of Gqα via RNAi employing a modified Rh1 containing a mCherry tag as the reporter. We show Rh1-mCherry is present in the rhabdomere of R1-6 photoreceptors in wild-type (Figure 4A) whereas Gqα-RNAi leads to the accumulation of internalized Rh1-mCherry and retinal degeneration as shown in 5-d old retinas (Figure 4B). This phenotype is similar to that observed in *norpA^P24^* (Figure 4 D), consistent with the notion that a reduced NorpA or Gqα activity greatly diminishes the Ca^2+^ influx critical for the recycling of the internalized rhodopsin (Alloway et al., 2000; Kiselev et al., 2000).

**Figure 4:**
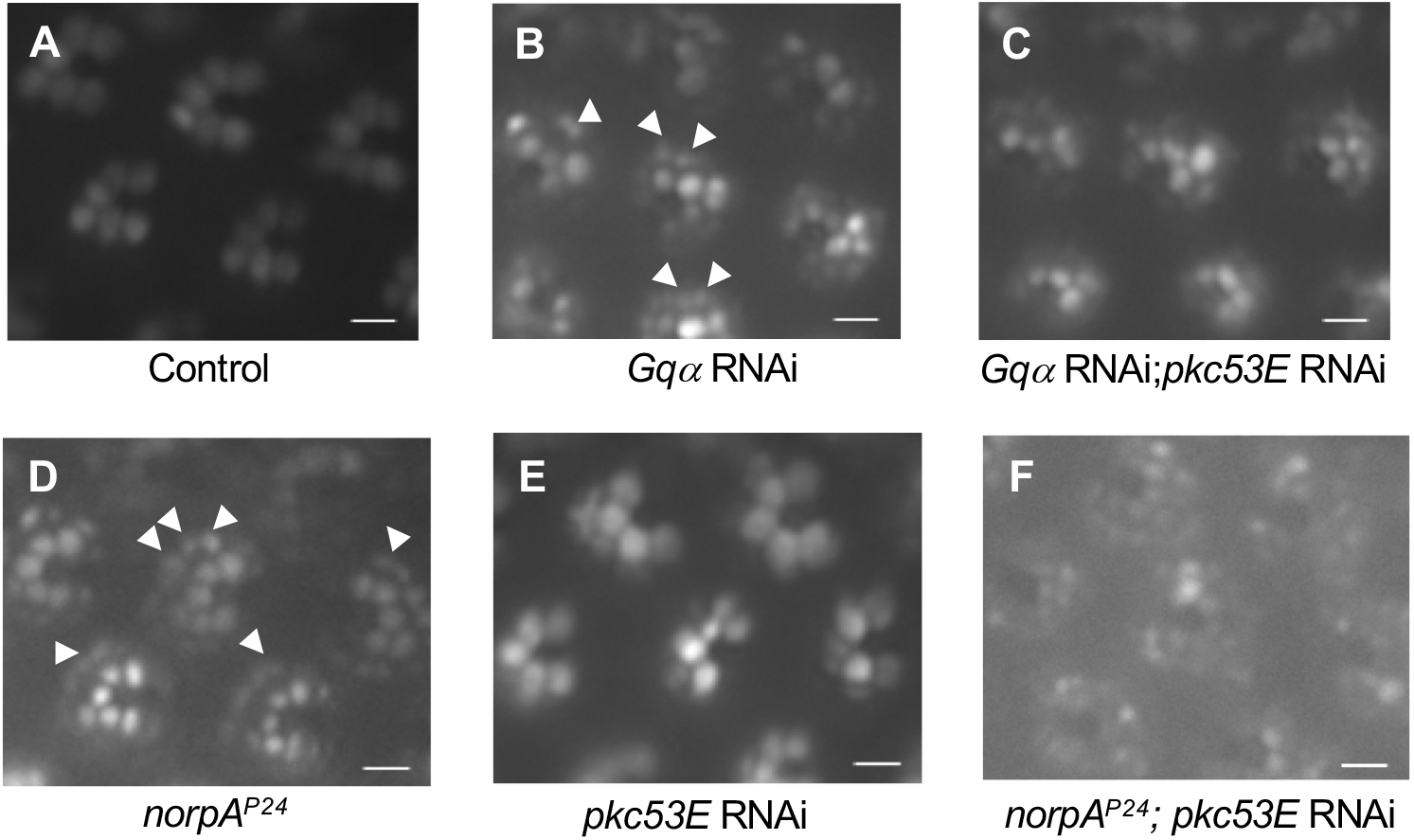
Pkc53E-RNAi does not modify degeneration caused by Gqα-RNAi. Gqα-RNAi leads to retinal degeneration that is accompanied by the accumulation of internalized Rh1 rhodopsin (**B**), similar to *norpA^P24^* (**D**) when compared to wild-type (**A**). Rh1-mCherry was used as the reporter. Internalized Rh1-mCherry are present in large vesicles surrounding the rhabdomere and some are marked with arrows. Retinal degeneration of Gqα-RNAi is not enhanced by Pkc53E-RNAi (**C**), unlike degeneration of *norpA^P24^* (**F**). Shown are retinas of 6-day old flies. Scale bars, 5 μm

We explored whether Gqα couples to Plc21C leading to the activation of Pkc53E in *norpA^P24^* photoreceptors. As mentioned before, in *norpA^P24^* photoreceptors Pkc53E remains active as degeneration is enhanced by Pkc53E-RNAi (Figure 4D, F). In contrast, Pkc53E-RNAi did not modify retinal degeneration caused by Gqα-RNAi (Figure 4B, C), strongly supporting that Pkc53E is not active when Gqα is greatly reduced. Similarly, eye-PKC RNAi did not modify retinal degeneration caused by Gqα-RNAi, consistent with the notion that DAG synthesis is blocked by Gqα-RNAi. Moreover, Gαq-RNAi did not modify retinal degeneration of *norpA^P24^* photoreceptors (not shown). Based on the findings, we propose that Gqα may couple to Plc21C leading to the activation of Pkc53E when NorpA is absent.

### Molecular characterization of *Drosophila* adducin

Members of the cPKC family have been implicated in remodeling of the actin cytoskeleton (Larsson, 2006). We explored whether Pkc53E regulates adducin, a cytoskeletal protein critical for the assembly of the membrane skeleton by regulating the interaction between the actin filament and the spectrin network (Matsuoka et al., 2000). In vertebrates, there are three adducin homologs, α-adducin (ADD1), β-adducin (ADD2), and γ-adducin (ADD3). Functional adducin consists of heterotetramers that are made up of either αβ or αγ heterodimer (Matsuoka et al., 2000).

*Drosophila* contains one adducin gene, *hts* (Yue and Spradling, 1992), which generates three adducin-like polypeptides of 739 (Add1; isoforms Q, R), 718 (Add2; isoforms I, L, M), and 716 aa (Add3; isoforms O, S) with different C-terminal sequences (Figure 5B). We show that mRNA transcripts corresponding to these three isoforms are expressed in photoreceptors (Figure 5C). Like mammalian isoforms, *Drosophila* adducin contains a globular domain at the N-terminus, a neck domain, and a C-terminal tail with the MARCKS-like domain that contains the conserved PKC phosphorylation sites (Figure 5A, B).

**Figure 5:**
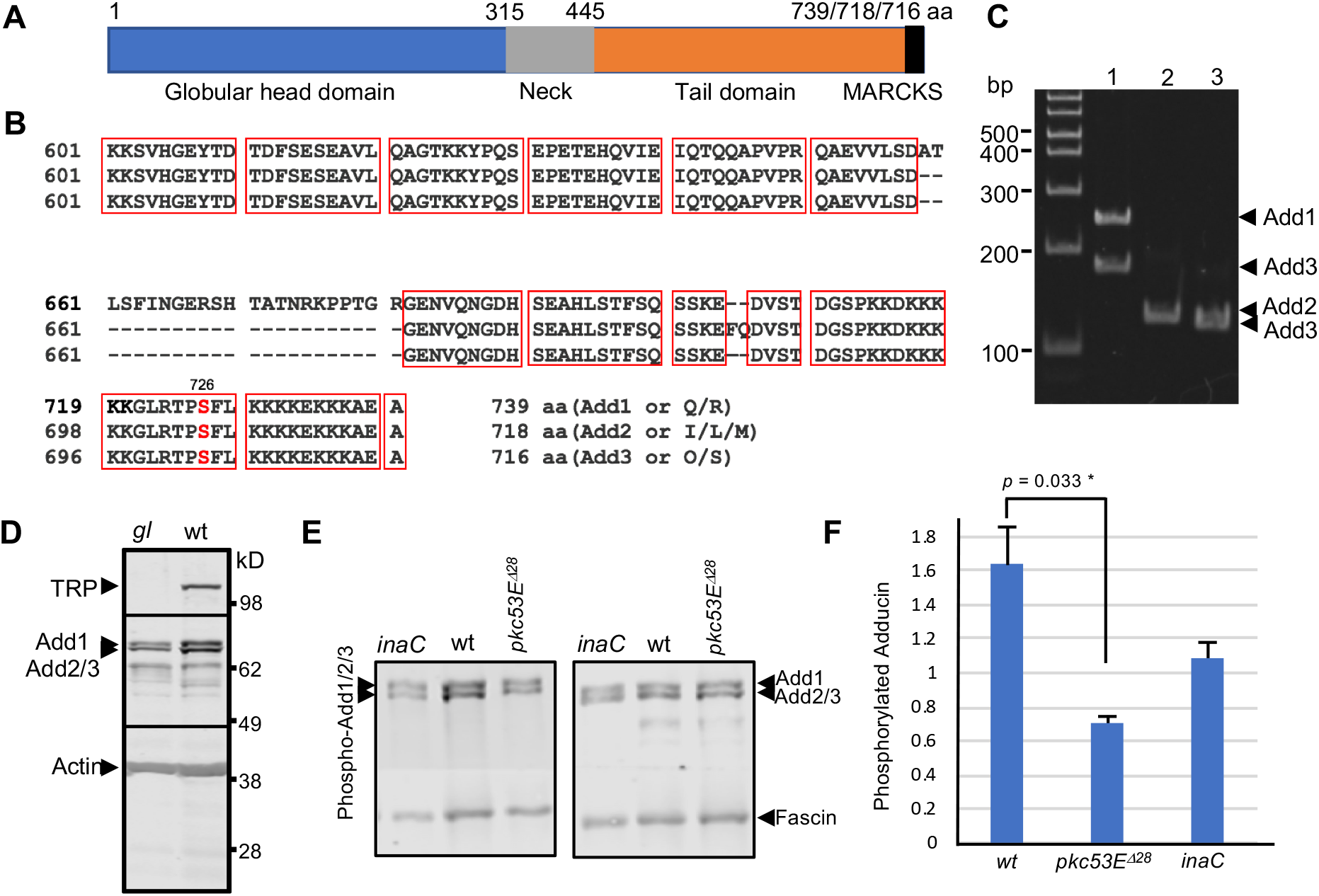
Expression of adducin-like isoforms of Hts and phosphorylation by Pkc53E. **A**, A graphic representation of the domain structure in three *Drosophila* adducin isoforms that consist of 739 (Add1), 718 (Add2), or 716 aa (Add3), respectively. All isoforms contain a globular head domain, a neck domain, and the variable C-terminal tail sequence with the MARCKS-related domain. **B**, The alignment of the C-terminal sequences that contain part of the tail domain. Amino acid sequence identities are boxed and the conserved PKC phosphorylation sites at Ser^726/705/703^ are highlighted in red. **C**, Expression of three adducin isoforms by RT/PCR. Sequence-specific primers (see Materials and Methods) were used for each isoform. The predicted PCR product is indicated on the right. DNA molecular weight standards are shown on the left. **D**, Western blot analysis of wild-type and *glass* head extracts using monoclonal antibodies for Hts (1B1), TRP, and actin, respectively. Both wild-type and *glass* head extracts contain Hts including two strong bands that may correspond to 739 aa (Add1) and 718/716 aa (Add2/Add3) isoforms. **E**, Phosphorylation of adducin at the conserved PKC site, Ser^726/705/703^ in each isoform. Shown are Western blots probed with either 1B1 (right panel) or polyclonal antibodies for phospho-Ser^726/705/703^ (left). Both adducin bands encompassing all isoforms were phosphorylated *in vivo*, and phosphorylation was greatly reduced in *Pkc53ED^Δ28^*, but not in *inaC^P209^*. Total and phosphorylated adducin were quantified using fascin as a loading control. Levels of phosphorylated adducin in the mutants were compared in a histogram (**F**). Each lane contained protein extracts prepared from four retinas.

We investigated the expression of adducin isoforms in photoreceptors by Western blotting using a monoclonal antibody (1B1) that recognizes its N-terminal sequence. We observed multiple protein bands including two prominent bands above 80 kD in size (Figure 5D), which may correspond to Add1 (739 aa, upper band), and Add2/Add3 (718/716 aa, lower band), respectively. Significantly, these protein isoforms appear ubiquitously expressed as they are present in head extracts from both wild-type and *glass* mutants (Figure 5D).

### Reduced phosphorylation of adducin at Ser^726/705/703^ in a *pkc53E* null allele

To explore whether Pkc53E phosphorylates adducin at the conserved PKC site, Ser^726/705/703^, in each isoform, *in vivo*, we employed Western blotting using polyclonal antibodies that recognize phosphorylated Ser^726/705/703^. Indeed, we observed that both upper and lower bands of adducin were phosphorylated in wild-type retinal extracts (Figure 5E). We calculated the steady-state level of total phosphorylated adducin, and demonstrate that it was reduced by 57.5 % ± 5.2% (n = 3) in *pkc53ED^Δ28^*, but was not significantly affected in *inaC^P209^* (Figure 5E, F). Together, our findings strongly support that phosphorylation of adducin is partly regulated by Pkc53E *in vivo*.

### Hts-RNAi leads to loss of rhabdomeres or ommatidial clusters similar to Pkc53E-RNAi

We explored the function of adducin in photoreceptors via Hts-RNAi. First, we validated Hts-RNAi using Western blotting and show that the level of adducin in the fly head was not greatly reduced by Hts-RNAi (Figure 6A), which may be due to the ubiquitous expression of adducin. Therefore, we chose a transgenic fly that co-expressed a mCherry tagged adducin (Add2-mCherry) that served as a marker for Hts-RNAi in photoreceptors (Figure 6A). Indeed, we demonstrate that the expression of the tagged adducin was greatly reduced by Hts-RNAi via Western blotting (Figure 6A). We presumed that endogenously expressed adducin could be also greatly reduced in photoreceptors.

**Figure 6:**
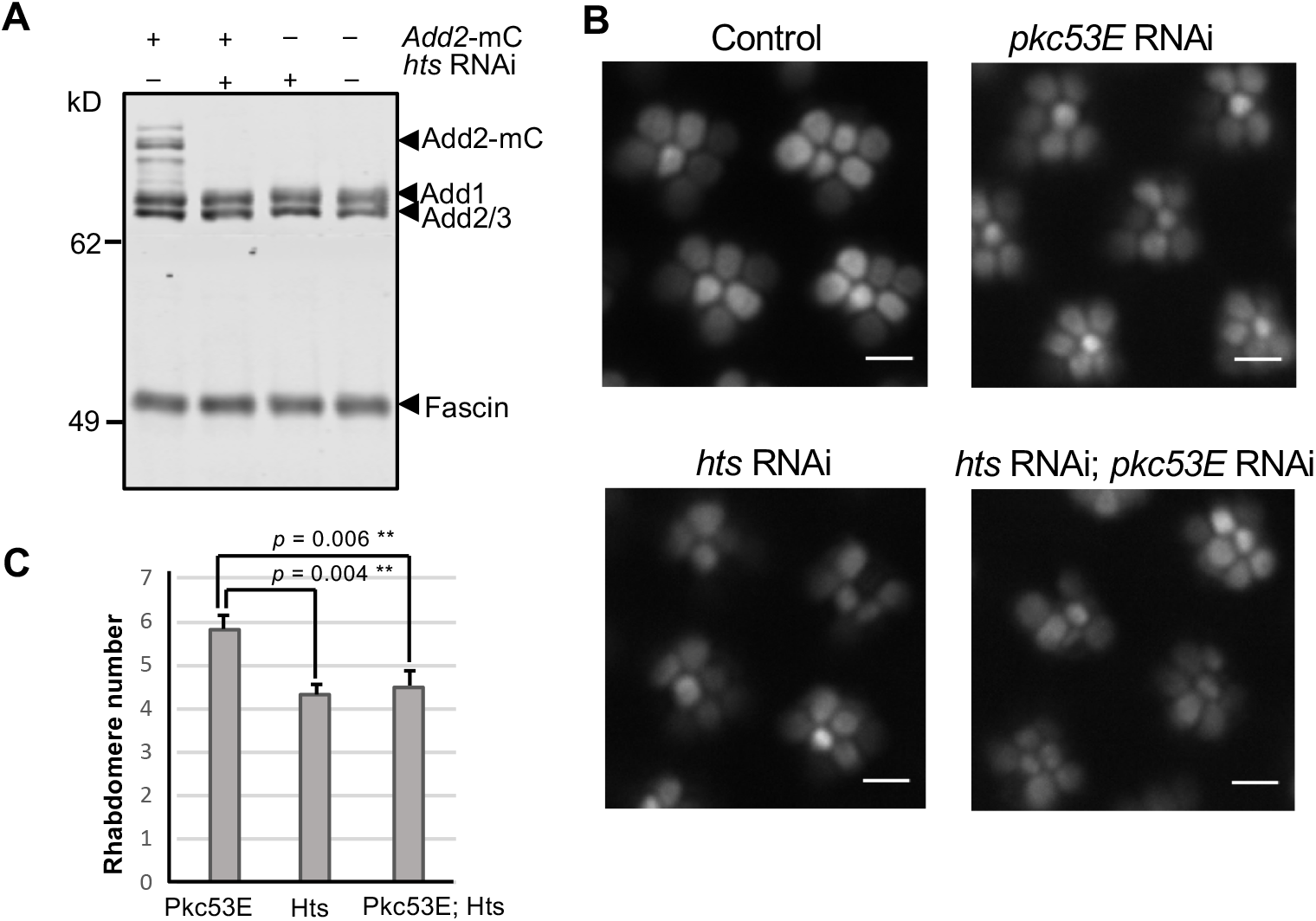
Hts-RNAi leads to a more severe retinal degeneration than Pkc53E-RNAi. **A**, Validation of Hts-RNAi by the loss of tagged Add2 expression via Western blotting. The mCherry tagged Add2 with an estimated size of 115 kD migrates above the endogenous adducin in SDS/PAGE (10%). Protein size markers are indicated on the left. **B**, Retinal morphology of Hts-RNAi using Actin-GFP as the marker. Hts-RNAi exhibits a phenotype with loss of rhabdomeres, which appears more severe than that of Pkc53E-RNAi. Shown are retinas of 7-day old flies. Scale bars, 5 μm. **C**, Quantitation of rhabdomere numbers in Pkc53E-RNAi, Hts-RNAi, and double RNAi (n = 9).

Using Actin-GFP as a reporter, we show that Hts-RNAi led to a loss of rhabdomeres and occasionally, missing ommatidia clusters, in 7-day old flies (Figure 6B). This defect is further exacerbated and a loss of rhabdomeres was observed in 14-day flies (not shown). This phenotype of Hts-RNAi is similar to that observed in Pkc53E-RNAi. Importantly, downregulation of both Pkc53E and adducin by RNAi results in a phenotype similar to that of Hts-RNA (Figure 6B, C), suggesting that adducin and Pkc53E are likely to act in the similar or same pathway involved in the maintenance of the actin cytoskeleton in photoreceptors.

### Overexpression of a tagged adducin leads to apical expansion of rhabdomeres

To further explore the role of adducin, we overexpressed in photoreceptors a modified adducin (Add2-mCherry). We show that overexpression led to sporadic ‘apical expansion’ characterized by the overgrowth of the actin-based cytoskeleton in the rhabdomere (Figure 7A). Importantly, this abnormal expansion occurs in all rhabdomeres within the same ommatidia, in which the diameter of rhabdomere increases about 2.73 ± 0.31 (n = 5) fold, compared to that of wild-type. Furthermore, the expansion affects the patterning of rhabdomere clusters as the typical trapezoidal arrangement of R1-6 rhabdomeres becomes less evident (Figure 7 A, right panel). The expansion phenotype is dependent on the copy number of the transgene; one copy resulted in expansion in 32.3 ± 13.5% (n = 10) while two copies gave rise to 46.1 ± 3.3% (n = 6) of abnormal ommatidia clusters. Despite the expansion, the external appearance of the compound eye is not significantly altered (not shown).

**Figure 7:**
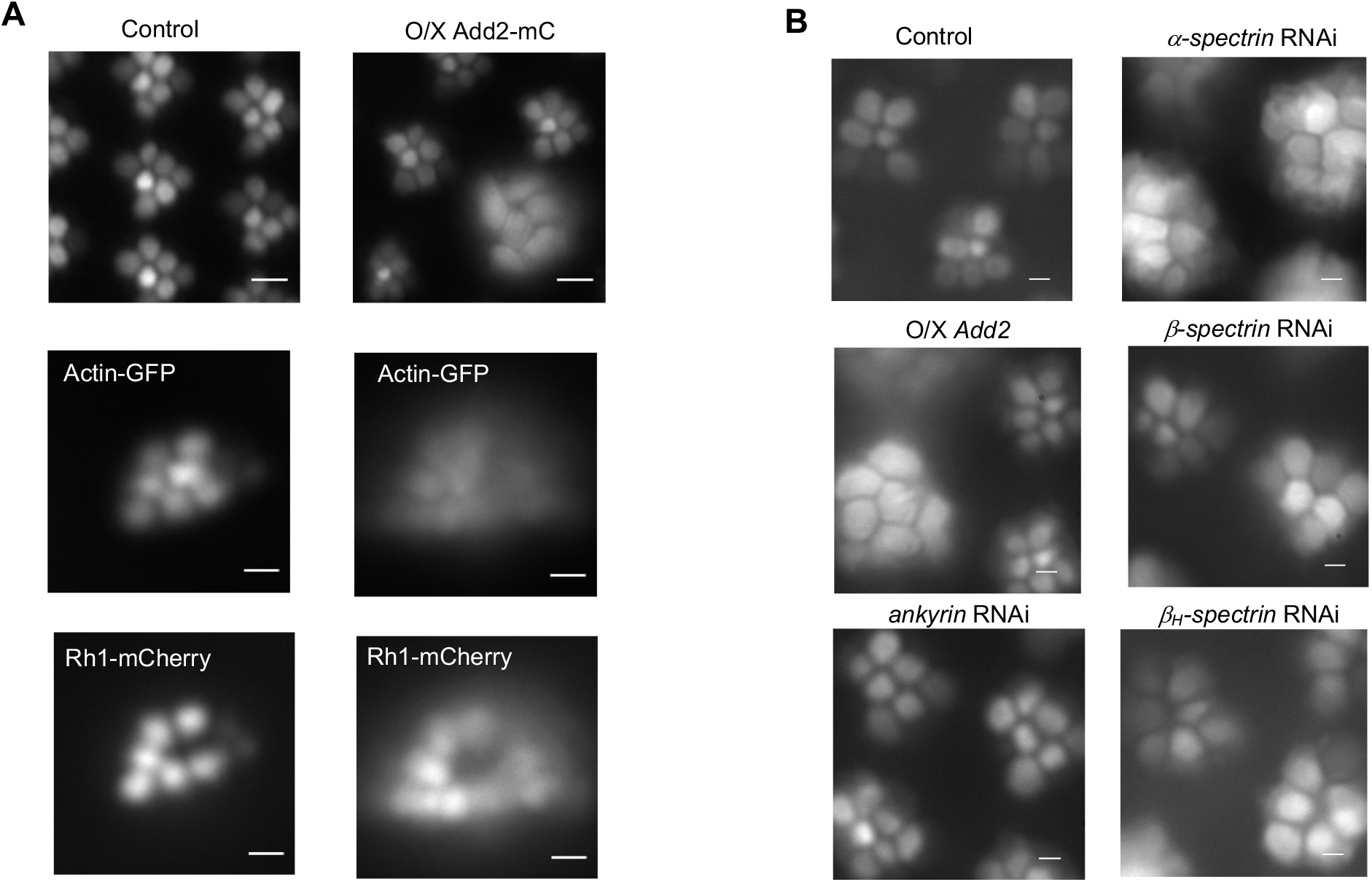
Overexpression of adducin leads to apical expansion of the actin cytoskeleton similar to β-spectrin-RNAi. **A**, Overexpression of mCherry tagged adducin, Add2, led to the apical expansion of rhabdomeres. In the compound eye, about 32.3 ± 13.5 % (n = 10) of ommatidia clusters (right panel, top) contain enlarged rhabdomeres that seem to project further outward when compared to control (left panel, top). The diameter of rhabdomere is increased to 2.73 ± 0.31 (n = 5) fold. The dpp becomes less distinct as visualized either by Actin-GFP (middle) or Rh1-mCherry (bottom), compared to wild-type (left panel). **B**, Spectrin-RNAi also leads to apical expansion. α-spectrin-RNAi (right panel, top) results in an enlarged actin cytoskeleton in all rhabdomeres. In contrast, β-spectrin-RNAi display enlarged rhabdomeres with reduced frequency compared to the expression of tagged adducin. β_H_-spectrin/karst-RNAi (right panel) also leads to apical expansion accompanied by the abnormal arrangement of rhabdomeres. In contrast, ankyrin1-RNAi has no significant effect on retinal morphology. Scale bars in **A**, 5 μm (top panel), 20 μm (bottom panels), and in **B**, 2 μm.

The presence of these expanded rhabdomeres also led to a distorted dpp (deep pseudopupil) (Franceschini, 1972) from either Actin-GFP or Rh1-mCherry (Figure 7A) as the alignment of the optical axis within the compound eye was affected (Figure 7A, middle, lower panels). This phenotype is detected in late pupal photoreceptors, indicating that it occurs during the morphogenesis of rhabdomeres following the differentiation of photoreceptors. We propose that adducin is critically involved in the regulation of rhabdomere expansion in pupal photoreceptors.

### β-Spectrin-RNAi also leads to apical expansion of ommatidia

We explored the cause of the expansion by comparing the phenotype to that from the knockdown of several known components involved in the actin-spectrin network including α-spectrin, β-spectrin, β_H_-spectrin (karst), and ankyrin1 (Bennett and Baines, 2001). Indeed, knockdown of either α- or β-spectrin also led to expansion similar to that observed in the overexpression of tagged Add2 (Figure 7B). Specifically, β-spectrin-RNAi resulted in the apical expansion of the ommatidia cluster with a frequency, 15.1± 2.1% (n = 7), which is about half of those observed in flies expressing the tagged Add2. α-spectrin-RNAi also led to enlarged rhabdomeres with reduced numbers of ommatidia clusters while β_H_-spectrin/karst-RNAi also exhibited abnormal rhabdomere organization within clusters. In contrast, the down regulation of ankyrin1 by RNAi has no significant effect (Figure 7B).

Based on the findings, we propose that the tagged Add2 acts in a dominant-negative manner to impact the assembly of the actin-spectrin network. This modified Add2 appears incapable of engaging with spectrin similar to phosphorylated adducin, and consequently, its overexpression interferes with the function of endogenously expressed adducin for interacting with β-spectrin.

### Epistasis analysis of Add2-mCherry and Pkc53E supports that Pkc53E is acting upstream of Add2

To further explore the relationship between adducin and Pkc53E, we carried out epistasis analysis to determine which gene is acting upstream if both are in the same pathway. We generated and characterized double mutants expressing the tagged Add2 in the background of altered Pkc53E activities. Importantly, neither down regulation by RNAi nor overexpression of Pkc53E-B affected the function of Add2-mCherry as double mutants displayed the apical expansion phenotype of the tagged Add2 at two days post eclosion (Figure 8A), supporting that Pkc53E is likely acting upstream of Add2 to regulate the expansion of the actin cytoskeleton during rhabdomere biogenesis.

**Figure 8:**
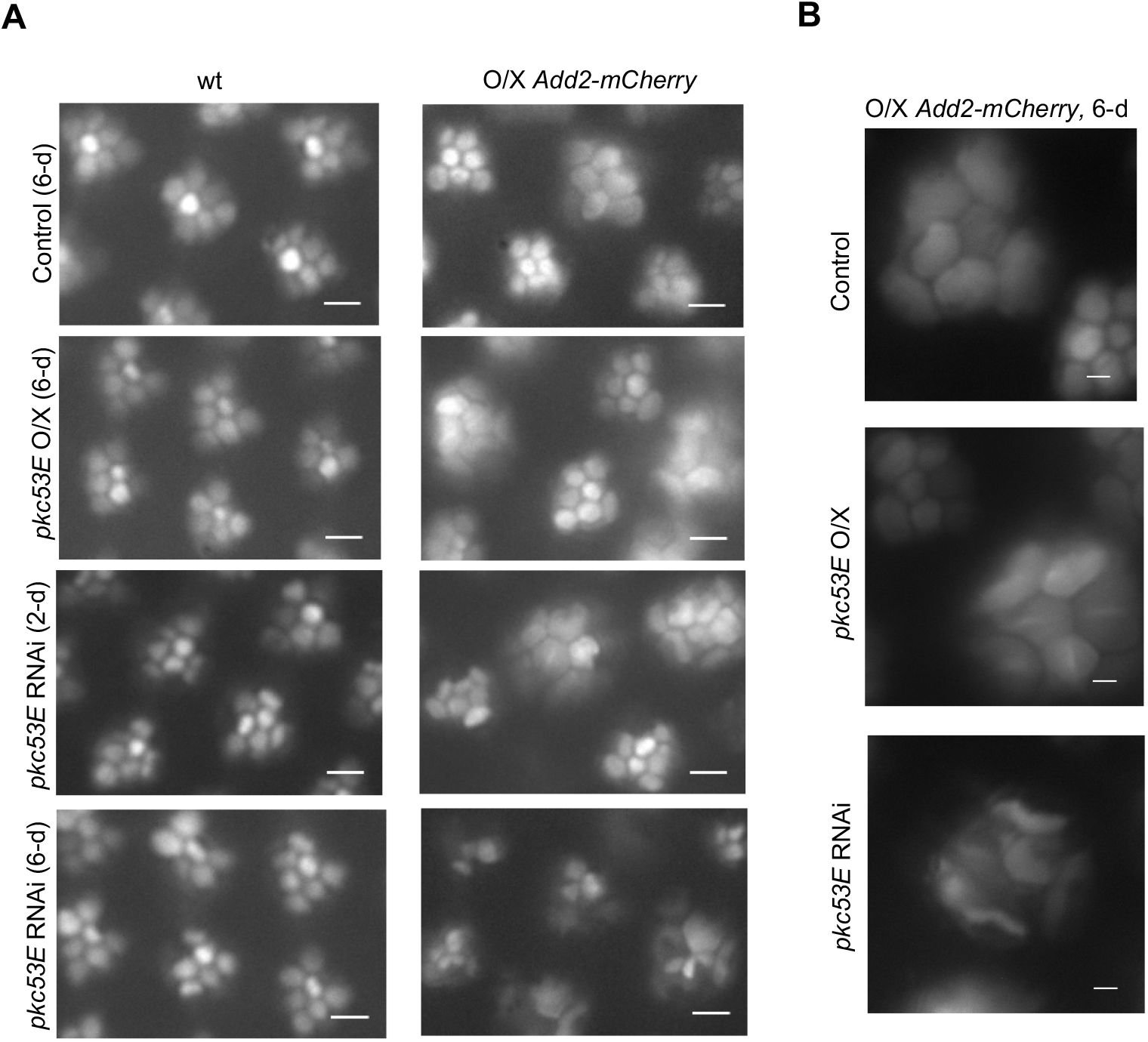
Double mutant analysis. **A**, Modulation of the apical expansion phenotype by Pkc53E. Effects of RNAi or overexpression of Pkc53E-B in wild-type (left panel) or the Add2-mCherry overexpressing background (right panel) using Actin-GFP as the reporter. The age of flies is indicated. In the Add2-mCherry background, down regulation or overexpression of Pkc53E did not affect the apical expansion as shown in 2-day old retinas. In contrast, expression of Add2-mCherry accelerates retinal degeneration of Pkc53E-RNAi (bottom panel). **B**, Higher magnification (1000X) of the expanded rhabdomeres in flies overexpressing Add2-mCherry with either overexpression or down regulation of Pkc53E. The abnormally enlarged rhabdomeres became distorted or flattened in adult photoreceptors when pKC53E is down regulated. Scale bars 5 μm in **A** and 2 μm in **B**.

In adult photoreceptors, overexpression of the Pkc53E-B did not affect the retinal morphology of flies expressing tagged Add2 (Figure 8A, B), further suggesting that Pkc53E is acting upstream of adducin. However, expression of the dominant-negative Add2 accelerated retinal degeneration of Pkc53E-RNAi as 6-day old double mutants displayed severe deterioration of the actin cytoskeleton. The degeneration phenotype includes missing or distorted rhabdomeres especially in those expanded and enlarged rhabdomeres (Figure 8A, B). A more severe phenotype of the double mutant may support that Pkc53E and adducin act in parallel pathways to regulate the rhabdomere of adult photoreceptors.

## Discussion

Here we explored the role of a photoreceptor-specific isoform of Pkc53E, Pkc53E-B, in the maintenance of the actin cytoskeleton of rhabdomeres in *Drosophila*. We show that Pkc53E-B is likely activated downstream of NorpA in the visual cascade. However, when NorpA is absent, Plc21C (Shortridge et al., 1991) may be coupled to Gq to switch on Pkc53E. We tested whether Pkc53E regulates adducin, a cytoskeletal protein critical for the assembly of the membrane skeleton (Baines, 2010). Adducin dynamically regulates the linkage between actin filaments and the spectrin network, which could be negatively affected by PKC phosphorylation (Barkalow et al., 2003; Matsuoka et al., 1998; Matsuoka et al., 2000). Indeed, in a null allele of *pkc53E*, we observed a reduced level of phosphorylated adducin. Furthermore, Hts-RNAi resulted in missing rhabdomeres, which is more severe than Pkc53E-RNAi, suggesting that both Pkc53E and adducin are acting in similar pathways in photoreceptors. Interestingly, overexpression of a mCherry tagged adducin leads to the apical expansion of rhabdomeres, which is also observed in β-spectrin-RNAi. We speculate that the expansion phenotype is caused by the interference of the actin-spectrin interaction that impacts rhabdomere morphogenesis during development.

Thus, in adult photoreceptors a rise of intracellular Ca^2+^ and the release of DAG accompanying the activation of PLCβ4 elevates the Pkc53E activity: Pkc53E phosphorylates adducin and releases it from the actin filament and the spectrin complex. Phosphorylated adducin may undergo dephosphorylation by cytosolic protein phosphatases to re-establish the link with the actin-spectrin network. The dynamic regulation of Pkc53E by light serves to fine-tune the linkage between the spectrin-based membrane skeleton and the plasma membrane, which may modulate the signaling in part by controlling protein localization and turnover.

### Pkc53E-B is a photoreceptor-specific isoform that regulates the actin cytoskeleton of rhabdomeres

Visual signaling leads to the activation of Pkc53E and eye-PKC. Eye-PKC is involved in fast activation and deactivation of the visual response (Hardie et al., 1993; Ranganathan et al., 1991; Smith et al., 1991) (Figure 9). We performed molecular characterization of Pkc53E and established that Pkc53E-B is highly expressed in photoreceptors. Consistently, Pkc53E-B mRNA is drastically reduced by Pkc53E-RNAi using the photoreceptor-specific GMR driver. Pkc53E-B is the longest protein isoform consisting of 679 aa that displays 57% sequence similarities to eye-PKC. We demonstrate, for the first time, that loss of function of *pkc53E* leads to retinal degeneration, supporting its critical role in the maintenance of photoreceptors. Pkc53E-B is likely acting downstream of NorpA/PLCβ4. However, we discovered that both Pkc53E and eye-PKC are partially active in *norpA^P24^* photoreceptors, suggesting that a partial loss of function of Pkc53E may contribute to the retinal degeneration in *norpA^P24^* photoreceptors.

**Figure 9:**
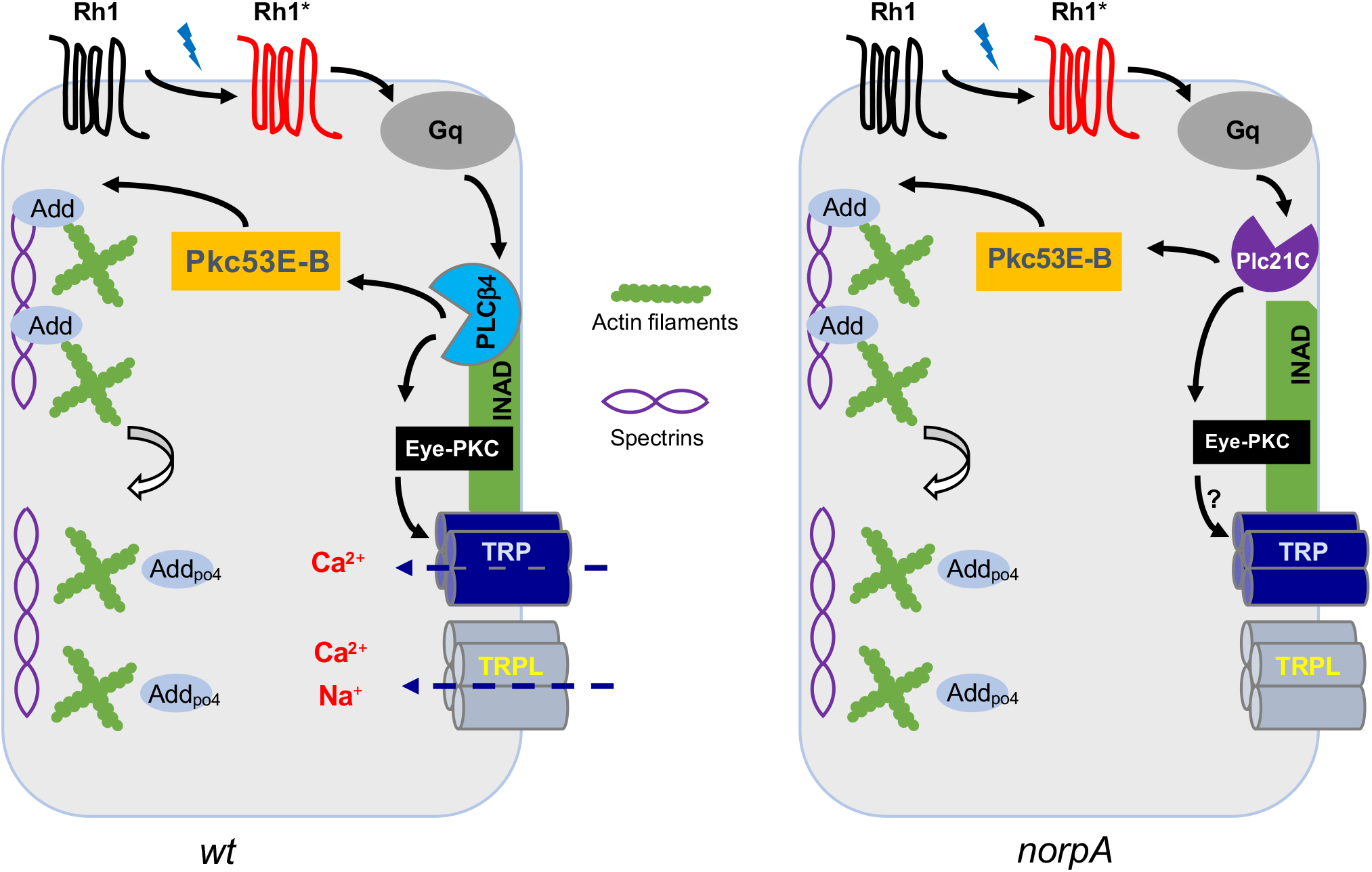
Role of Pkc53E to regulate adducin to promote the maintenance of the actin cytoskeleton. In wild-type photoreceptors (left panel), light activates rhodopsin (Rh1) that couples to Gq leading to the activation of PLCβ4 /NorpA. PLCβ4 generates DAG and increases Ca^2+^ leading to the activation of eye-PKC and Pkc53E. Eye-PKC phosphorylates the TRP Ca^2+^ channel to promote fast deactivation of the visual response. In contrast, Pkc53E phosphorylates adducin (Add) to orchestrate the remodeling of the actin cytoskeleton by uncoupling the interaction between actin filaments and the spectrin network. In *norpA^P24^* photoreceptors (right panel), Plc21C is activated instead of PLCβ4 leading to the activation of eye-PKC and Pkc53E.

We investigated the NorpA-independent activation of Pkc53E, and propose an alternate mechanism in which Plc21C is deployed leading to the activation of two PKC’s (Figure 9). It is likely that a loss of NorpA leads to unregulated Gqα, as NorpA serves as GTPase activating protein for inactivating Gqα (Cook et al., 2000). The prolonged Gq activity thus may promote its coupling to Plc21C.

### The function of adducin and its regulation of the spectrin network

The *Drosophila* adducin gene (*hts*) was originally identified for its role in the ring canal formation of the oocyte (Yue and Spradling, 1992). The *hts* locus encodes at least eight polypeptides including three adducin-like proteins containing the MARCKS-related domain (Flybase). These three isoforms vary in their C-terminal sequences and may correspond to the three mammalian homologs (Citterio et al., 1999; Joshi et al., 1991). Previously, a doublet of adducin-like polypeptides was observed in the Western blot (Pielage et al., 2011; Wang et al., 2014). Here we show that the adducin doublet is likely to encompass three related polypeptides. All adducin isoforms appear phosphorylated in photoreceptors.

We overexpressed a mCherry tagged Add2 in photoreceptors and observed the apical expansion characterized by the overgrowth of the actin cytoskeleton of the rhabdomere in about 32% of ommatidia clusters. This expansion phenotype is similar to that observed in β-spectrin-RNAi, suggesting that expression of the modified Add2 interferes with the formation of the spectrin network. Spectrins (α-spectrin, β-spectrin, and β_H_-spectrin) form oligomers that are the basis of the spectrin-based membrane skeleton, which is localized in the cytoplasmic side of the plasma membrane (Bennett and Baines, 2001). Specifically, α/β_H_-spectrins are localized at the apical domain (Pellikka et al., 2002) while α/β-spectrins in the basolateral domain of photoreceptors (Chen et al., 2009). It has been shown that removal of either α- or β-spectrin leads to the expanded apical domain while overexpression of β-spectrin reduces it (Chen et al., 2009), suggesting that basolaterally localized α/β-spectrins are critical for the maintenance of the apical domain. Moreover, spectrins may transmit the intracellular actomyosin force to the cell membrane to regulate cell shape as a loss of spectrins led to the overgrowth of cells (Deng et al., 2020; Fletcher et al., 2015). How each mechanism contributes to the rhabdomere expansion remains to be explored.

While adducin was first identified for its role in the assembly of the spectrin-based membrane cytoskeleton in erythrocytes (Matsuoka et al., 2000), it appears to exert diverse functions. In epithelia, adducin is critical for cellcell adhesion (Abdi and Bennett, 2008), regulating cell junctions and permeability (Naydenov and Ivanov, 2011). Interestingly, adducin was shown to be localized in the nucleus of cultured cells that were not forming contact with neighboring cells (Chen et al., 2011). In the nervous system, adducin was shown to be involved in the assembly of synapses by orchestrating the remodeling of dendritic spines (Bednarek and Caroni, 2011; Engmann et al., 2016); loss of β-adducin reduces spine formation affecting synaptic plasticity (Rabenstein et al., 2005).

To sum up, adducin regulates the interaction between actin filaments and the spectrin network. Here we uncovered two distinct functions of adducin in *Drosophila*. In adult photoreceptors, adducin is involved in the maintenance of the actin cytoskeleton supporting the rhabdomere; during pupal development, adducin is critical for rhabdomere morphogenesis.

### Relationship between adducin and Pkc53E for the regulation of the membrane skeleton

It is important to note that the Pkc53E activity in photoreceptors is regulated by light to exert its negative impact on adducin (Figure 9). A rhythmic adducin activity allows periodic re-alignment between the plasma membrane and the actin cytoskeleton in response to the visual signaling. We propose that Pkc53E phosphorylates adducin in photoreceptors based on the following findings. We show that Hts-RNAi results in a severe loss of rhabdomeres, which is not modified by Pkc53E-RNAi, suggesting that both are likely acting in the same or similar pathway. Moreover, epistasis analysis using the dominant-negative Add2 showed that downregulation of Pkc53E did not modify the apical expansion of the tagged Add2, strongly suggesting that adducin is acting downstream of Pkc53E. Consistently, levels of phosphorylated adducin are greatly reduced in a null allele of *pkc53E*. Together, we hypothesize that Pkc53E may phosphorylate adducin following the transmembrane signaling. However, dominant-negative Add2 greatly enhanced degeneration of Pkc53E-RNAi, which appears to support that each protein participates in different pathways to regulate the cytoskeleton.

The well-characterized functions of adducin include (1) capping the growing end of the actin filament, (2) linking actin filaments to the spectrin network, and (3) bundling actin filaments (Matsuoka et al., 2000). How does overexpression of tagged Add2 act dominant-negatively to affect rhabdomere morphogenesis? It is likely that the inclusion of the mCherry tag negatively impacts its association with either spectrin and/or actin filaments. It is also possible that modified Add2 becomes constitutively phosphorylated by a protein kinase thereby acting like a phosphomimetic adducin that is incapable of the spectrin interaction. The function of adducin is likely regulated at multiple levels, which remain to be explored.

### A hypothetical model linking activation of Pkc53E to the reorganization of the actin cytoskeleton

In summary, Pkc53E-B plays a role in regulating the stability of the actin cytoskeleton. Pkc53E-B may phosphorylate adducin that dynamically controls the linkage between actin filaments and the spectrin network. Thus, transmembrane signaling that activates PLCβ or PLCγ leading to the activation of cPKC may transiently uncouple the spectrin network from actin filaments. This triggers the extension of actin filaments to promote the remodeling of the actin cytoskeleton, which is critical for maintaining the cell shape and possibly for regulating protein turnover in the plasma membrane. Remodeling of the membrane cytoskeleton may facilitate endocytosis and trafficking of transmembrane proteins thereby fine-tuning the signaling event. In the visual signaling in which activation of rhodopsin leads to an increase of cytosolic Ca^2+^ and transient activation of cPKC. Both Ca^2+^ and cPKC are involved in the stability of the actin cytoskeleton (Figure 9). Indeed, when the canonical pathway to turn on cPKC is missing, an alternate pathway that uses Plc21C may be deployed to support the maintenance of *Drosophila* photoreceptors (Figure 9). We propose that regulation of the actin cytoskeleton is a fundamental event for the PLC-mediated signaling mechanisms.

## Abbreviations

Aa: amino acids
Arr2: arrestin 2
cPKC: conventional/classical protein kinase C
DAG: diacylglycerol
dpp: deep pseudopupil
hts: hu-li tai shao
inaC: inactivation-no-afterpotential C
INAD: inactivation-no-afterpotential D
MARCKS: myristoylated alanine-rich C-kinase substrate
NorpA: no-receptor potential A
PKC: protein kinase C
PLC: phospholipase C
PLD: phospholipase D
RNAi: RNA interference
RT/PCR: reverse transcription/polymerase chain reaction
TRP: transient receptor potential
TRPL: TRP-like

