## Supplemental Figure 1 for "Transmembrane Signaling Regulates the Remodeling of the Actin Cytoskeleton: Roles of PKC and Adducin"

**Figure S1. Characterization of a Pkc53E null allele, and validation of Gαq-RNAi, and eye-PKC-RNAi by Western blot analysis**

**A**, The RT/PCR analysis of wild-type and the *pkc53E* mutant. The PCR products corresponding to rp49 and Pkc53E isoforms are indicated on the right The *pkc53E* mRNA expression is detected in wild-type but missing in the null allele. **B,** Gαq-RNAi leads to a greatly reduced level of Gαq while the eye-PKC level is drastically decreased by eye-PKC-RNAi (**C**). INAD and TRP were used for loading controls.

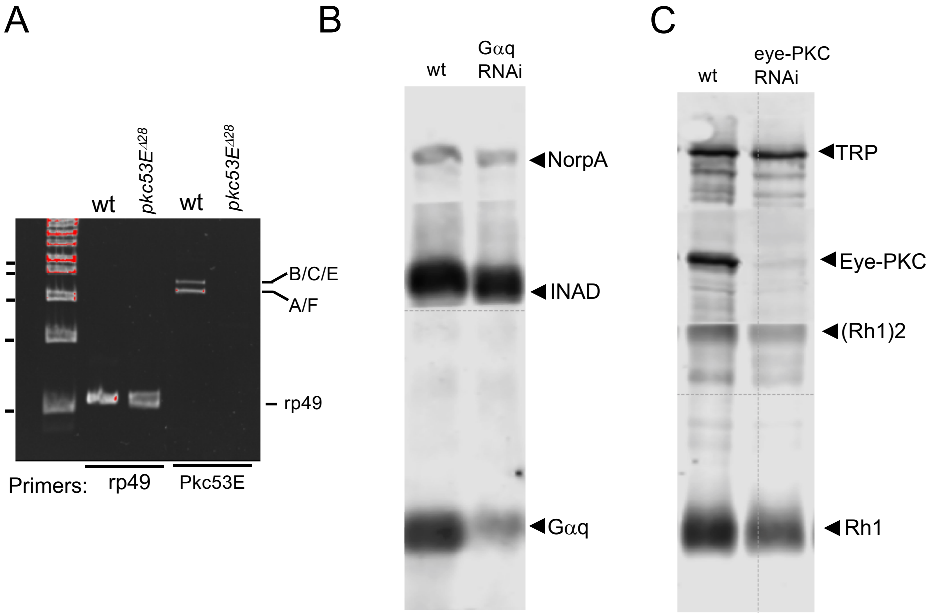
A
wt
pkc53E$^{\Delta 28}$
wt
pkc53E$^{\Delta 28}$
B/C/E
A/F
rp49
Primers:
rp49
Pkc53E
B
wt
Gαq
RNAi
NorpA
INAD
Gαq
C
wt
eye-PKC
RNAi
TRP
Eye-PKC
(Rh1)2
Rh1

Figure S1
